# Farmed cricket performance remains stable over five generations of rearing on a waste-based diet

**DOI:** 10.1101/2025.06.24.660593

**Authors:** S.Y. Kasdorf, S.M. Bertram, H.A. MacMillan

## Abstract

Farmed insects like crickets offer a sustainable protein source to feed the growing global population. A benefit of cricket farming is the potential to use waste diets instead of unsustainable, expensive feeds. Brewer’s spent grain is a nutritionally valuable organic waste product that has been used to rear crickets in single generation studies. However, the long-term effects of spent grain-based feeds are unclear, which makes incorporation into commercial feed risky for producers. We reared a farmed cricket (*Gryllodes sigillatus*) for five generations on a high inclusion (75%) spent grain diet. Crickets reared on spent grain were 18.5% smaller at adulthood than control crickets reared on farm feed, resulting in decreased yield (mass of crickets harvested), but were able to reproduce and had high survival rates. Cricket performance remained stable over five generations, indicating that spent grain contains adequate nutrition to support long-term cricket production. We also reared crickets on a gradual inclusion spent grain diet that increased from 15-75% spent grain over five generations. While this “weaning” approach did not improve cricket performance on high-inclusion spent grain diets, crickets on low inclusion diets (15-30%) displayed nearly a 30% increase in survival and yield compared to those fed the control. Therefore, inclusion of low amounts of spent grain in cricket feed may not only be beneficial from an environmental and feed cost perspective, but also from a production yield perspective. Our findings are the first to show that spent grain is a suitable feed ingredient for long-term rearing of farmed crickets.

## Introduction

Farmed insects are gaining popularity as a source of sustainable food and feed, leading to a growing market for insect production in North America (Larouche *et al*., 2023). Production success of the farmed insect industry is heavily dependent on the quantity of insect biomass produced (yield). Yield is influenced by insect life history traits like survival (number of insects available to harvest), body mass (weight of insects harvested), development time (time needed to reach harvestable size) and fecundity (ability to produce the next generation) (Kong *et al*., 2025; Muzzatti *et al*., 2024). Insect life history is shaped by independent and interactive effects of many external factors including diet (Muzzatti *et al*., 2024), temperature (Eberle *et al*., 2022) and rearing density (Mahavidanage *et al*., 2023). While ideal production conditions would optimize all of these traits, finding trait optima requires close collaboration between industry and academia, connecting fundamental research on insect physiology and ecology with the needs of producers (Kong *et al*., 2024; Larouche *et al*., 2023; Tomberlin *et al*., 2022).

Feed ingredient choice is a particularly important consideration for insect producers due to a need to balance nutritional value and feed cost. Constraints on nutrient availability in feed relative to insect nutritional requirements lead to a reduction in productivity. For example, insufficient dietary protein may limit nitrogen availability to allocate towards egg production, resulting in lower egg counts (Boggs, 2009). Many producers rely on traditional, grain-based feeds similar to those used for vertebrate livestock that have a balanced nutritional profile capable of supporting high rates of insect growth and survival (Lundy and Parrella, 2015; Oonincx *et al*., 2015; Sorjonen *et al*., 2019). However, traditional feed ingredients like corn, soy and fishmeal are expensive (Donohue and Cunningham, 2009; Macusi *et al*., 2023), and this is an impediment to the economic viability of the edible insect industry (Biteau *et al*., 2024; Halloran *et al*., 2017). In addition, feed crops contribute significantly towards the environmental impact of livestock production, and, combined with pastures, use 40% of arable land on earth (Mottet *et al*., 2017). Therefore, there is a need to reduce both the economic and environmental cost of insect feed to support industry growth and global sustainability, while maintaining production success.

High-volume organic waste products like brewer’s spent grain are potential alternatives to traditional feed for insect rearing. The brewing industry produces over 36.4 million tonnes of spent grain worldwide each year (Zeko-Pivač *et al*., 2022), which is composed mainly of spent barley grain containing 15-24% protein (Mussatto *et al*., 2006; Rachwał *et al*., 2020; Thiago *et al*., 2014). Therefore, this product is both a valuable source of nutrition and relatively ubiquitous. Multiple farmed insect species have been raised on diets comprised fully or partially of spent grain, including several species of farmed cricket (Jucker *et al*., 2022; Kasdorf *et al*., 2025; Oonincx *et al*., 2015; Orinda *et al*., 2017; Sorjonen *et al*., 2019), black soldier flies (*Hermetia illlucens*) (Bava *et al*., 2019; Resconi *et al*., 2024) and mealworms (van Broekhoven *et al*., 2015). In addition to providing a cost-effective feed ingredient for producers, recovery of spent grain for commercial insect production provides a key opportunity to contribute towards establishment of a circular economy (CE). The CE framework aims to minimize resource consumption and waste output to support sustainable agriculture (Velasco-Muñoz *et al*., 2021), and the framework is mentioned as a potential benefit of insect farming due to insects’ capacity to convert waste (Madau *et al*., 2020). However, spent grain inclusion in insect diets can lead reductions in growth (Kasdorf *et al*., 2025; Lundy and Parrella, 2015; Sorjonen *et al*., 2019), and the advantages and disadvantages of spent grain diets need to be deeply understood before producers are likely to include this product in their feed.

To date, studies investigating the effects of spent grain inclusion in mass-rearing diets are almost exclusively performed on a single generation of insects. A key knowledge gap is whether or not farmed insects will respond differently to these novel diets over the long-term. By omitting measurements of important factors like reproductive success or changes in life history traits (i.e., performance) over time, these studies cannot reveal multigenerational benefits or costs of waste inclusion. Long-term declines in insect performance may be experienced due to a compounding effect of malnutrition or epigenetic factors. For example, spruce budworm (*Choristoneura fumiferana*) fed a low quality diet for four generations experienced negative impacts on life history like increased mortality and reduced female reproductive output that accumulated over time (Frago and Bauce, 2014). Conversely, black soldier fly larvae (*Hermetia illucens*) reared on a poor diet (wheat bran) displayed an increase in performance traits like larval mass after twelve generations (Gligorescu *et al*., 2023). As in these examples, a deeper understanding of spent-grain effects on insect life history would assist producers in making informed decisions about the potential risks and benefits of integrating waste-based diets into large-scale commercial farming. Here, we investigate the impacts of multigenerational rearing of banded crickets (*Gryllodes sigillatus*), a popular species for commercial production (Zielińska *et al*., 2015), on spent-grain-based diets. We maintained crickets for five generations on one of three experimental diets: a control (existing farm feed), a high inclusion diet (75% spent grain), and a gradual inclusion diet (increased incrementally from 15-75% spent grain over the course of the five generations). The purpose of the gradual inclusion diet was to investigate a “weaning” approach of gradually introducing a novel diet through multiple intermediary diets that has been used to successfully facilitate the transition of fish to formulated diets in aquaculture (Gisbert *et al*., 2016; Kubitza and Lovshin, 1999; Ljubobratović *et al*., 2015), although typically in a single generation.

## Methods

### Diet Treatments

Brewer’s spent grain, free of hops to avoid a bitter taste, was generously provided by Stray Dog Brewing Company in Ottawa, Ontario, Canada. The spent grain was dried in a drying oven (31574, Precision Scientific Co, Chicago, USA) at 30°C for approximately ten days, then ground using a Vevor® Commercial Grinding Machine and stored at -20°C. A control farm feed was obtained from Entomo Farms in Norwood, Ontario, Canada. All components of the feed were obtained dried and ground from Campbellford Farm Supply in Campbellford, Ontario, Canada. Five experimental diets were created in which the control feed was replaced by mass with spent grain at a proportion of 15%, 30%, 45%, 60% or 75% (Table 1). Micronutrient supplements (Hog Grower Premix, salt, CaCO_3_, phosphate) used in the farm feed were kept at a consistent proportion in all diets.

**Table 1.**
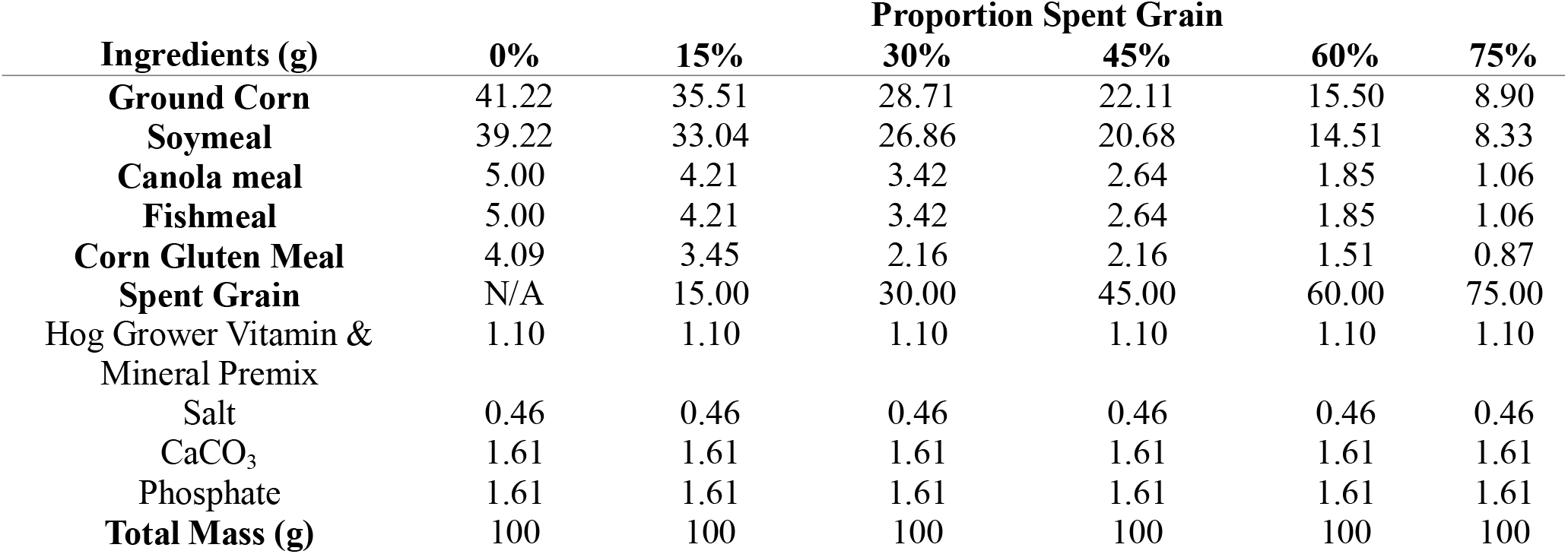
Ingredient composition (g) of diets.

| Ingredients (g) | Proportion Spent Grain |  |  |  |  |  |
| --- | --- | --- | --- | --- | --- | --- |
|  | 0% | 15% | 30% | 45% | 60% | 75% |
| **Ground Corn** | 41.22 | 35.51 | 28.71 | 22.11 | 15.50 | 8.90 |
| **Soymeal** | 39.22 | 33.04 | 26.86 | 20.68 | 14.51 | 8.33 |
| **Canola meal** | 5.00 | 4.21 | 3.42 | 2.64 | 1.85 | 1.06 |
| **Fishmeal** | 5.00 | 4.21 | 3.42 | 2.64 | 1.85 | 1.06 |
| **Corn Gluten Meal** | 4.09 | 3.45 | 2.16 | 2.16 | 1.51 | 0.87 |
| **Spent Grain** | N/A | 15.00 | 30.00 | 45.00 | 60.00 | 75.00 |
| Hog Grower Vitamin & Mineral Premix | 1.10 | 1.10 | 1.10 | 1.10 | 1.10 | 1.10 |
| Salt | 0.46 | 0.46 | 0.46 | 0.46 | 0.46 | 0.46 |
| CaCO<sub>3</sub> | 1.61 | 1.61 | 1.61 | 1.61 | 1.61 | 1.61 |
| Phosphate | 1.61 | 1.61 | 1.61 | 1.61 | 1.61 | 1.61 |
| **Total Mass (g)** | 100 | 100 | 100 | 100 | 100 | 100 |

### Rearing

Newly hatched crickets (*Gryllodes sigillatus*) were obtained from a stock population maintained at Carleton University that originated from Entomo Farms, Norwood, Canada. Hatchlings were randomly allocated to one of three diet treatments: a Control, High Inclusion (75%) Spent Grain or Gradual Inclusion (15-75%) Spent Grain (Table S1). Each diet was provided to crickets in five replicate plastic bins (49.5 × 38.1 × 35.2 cm) containing egg carton for shelter, a plastic vial of water plugged with cotton, and mesh in the lid for aeration (n ≈ 150 hatchlings per bin, estimated by mass). Crickets were allowed to feed *ab libitum* and all bins were kept in a greenhouse at 30°C, approximately 25% humidity and a 14L:10D photoperiod generated using LEDs for the duration of the experiment.

The experiment was run for five generations. Ten days after the first adult was observed in the majority of bins on a treatment, crickets were provided with damp peat moss for oviposition. After two-days of oviposition, the peat moss was removed to allow egg incubation (30°C, 25% humidity, 14L:10D) and hatchling emergence, and adult crickets were harvested (culled). A subset of the hatchlings was returned to the parental bin to restart the cycle (n ≈ 150 hatchlings per bin, estimated by mass).

### Data Collection

To monitor changes in life history traits over time, we quantified cricket adult body mass, adult body size, time to adulthood, survival rate, harvest yield, hatching yield and hatchling mass for each generation. At the end of every generation immediately prior to harvest, we obtained adult body mass and body size measurements (n = 10 crickets per bin, pseudorandomly selected). Mass was obtained using an analytical balance (AB135-S, Mettler Toledo, Columbus, USA), and body size was measured by capturing an image of each cricket using a microscope (Stemi 508, ZEISS, Oberkochen, Germany) with a colour camera (Axiocam 208, ZEISS) and ZEN Lite (ZEISS) software. Head width, pronotum width and pronotum length measurements were obtained from the images using ImageJ v.1.53 software (National Institutes of Health, Bethesda, MD, U.S.A.). Time to first adult emergence (by bin) was used as a proxy for development time and was recorded as the number of days between the date that hatchlings were added to a bin and the date that adults were first observed. Survival rate (by bin) was quantified by counting the number of living adult crickets in each bin at the end of every generation and dividing it by the starting number (estimated hatchlings by weight). Harvest yield (by bin) was defined as the total mass of crickets produced at the end of every generation, and was quantified by weighing all crickets as a group using an analytical balance (PA224 Pioneer™ Plus, OHAUS, Parisippany, USA). Hatchling yield (by bin) was defined as the total mass of hatchlings that emerged from peat moss (egg-laying substrate) within two days of hatching start, and was quantified by weighing all hatchings as a group using an analytical balance (AB135-S, Mettler Toledo, Columbus, USA). We obtained hatchling body mass measurements (n = random subset of 10 hatchlings per bin) at the start of every generation using the same balance.

### Statistical Analyses

All statistical analyses were performed using R (v4.2.3; (R Core Team, 2024)) software. Principal component analyses were used to obtain one value representative of overall body size using the three body size measurements (head width, pronotum width, pronotum length) obtained for the adult crickets (Visanuvimol and Bertram, 2011). The PC1 size value accounted for 93.4% (eigenvalue = 2.80) of total variance and was positively correlated with head width (loading score = 0.577), pronotum width (loading score = 0.584) and pronotum length (loading score = 0.570), indicating that it is a good representation of cricket size. We tested for normality using a Shapiro-Wilks normality test for data grouped by diet, generation and sex (adult body mass, adult body PC1 size), or diet and generation (survival, harvest yield, time to adulthood, hatchling mass and hatchling yield). In cases where crickets emerged on the same day, the normality test was not performed. Linear mixed effects models were used to determine the effect of diet, generation and sex on adult cricket body mass and body size (e.g., Cricket Body Mass ∼ Diet x Sex x Generation + (1 | Bin)) using replicate bin as a random effect. Linear mixed effects models were used to determine the effect of diet and generation on survival rate, harvest yield, time to adulthood, hatchling mass and hatchling yield (e.g., Survival Rate ∼ Diet x Generation + (1 | Bin)). Model output is shown in Table 2. To clarify differences among treatment groups, pairwise comparisons were performed for diet treatments at each generation and adjusted for multiple comparisons using Bonferonni correction.

**Table 2.**
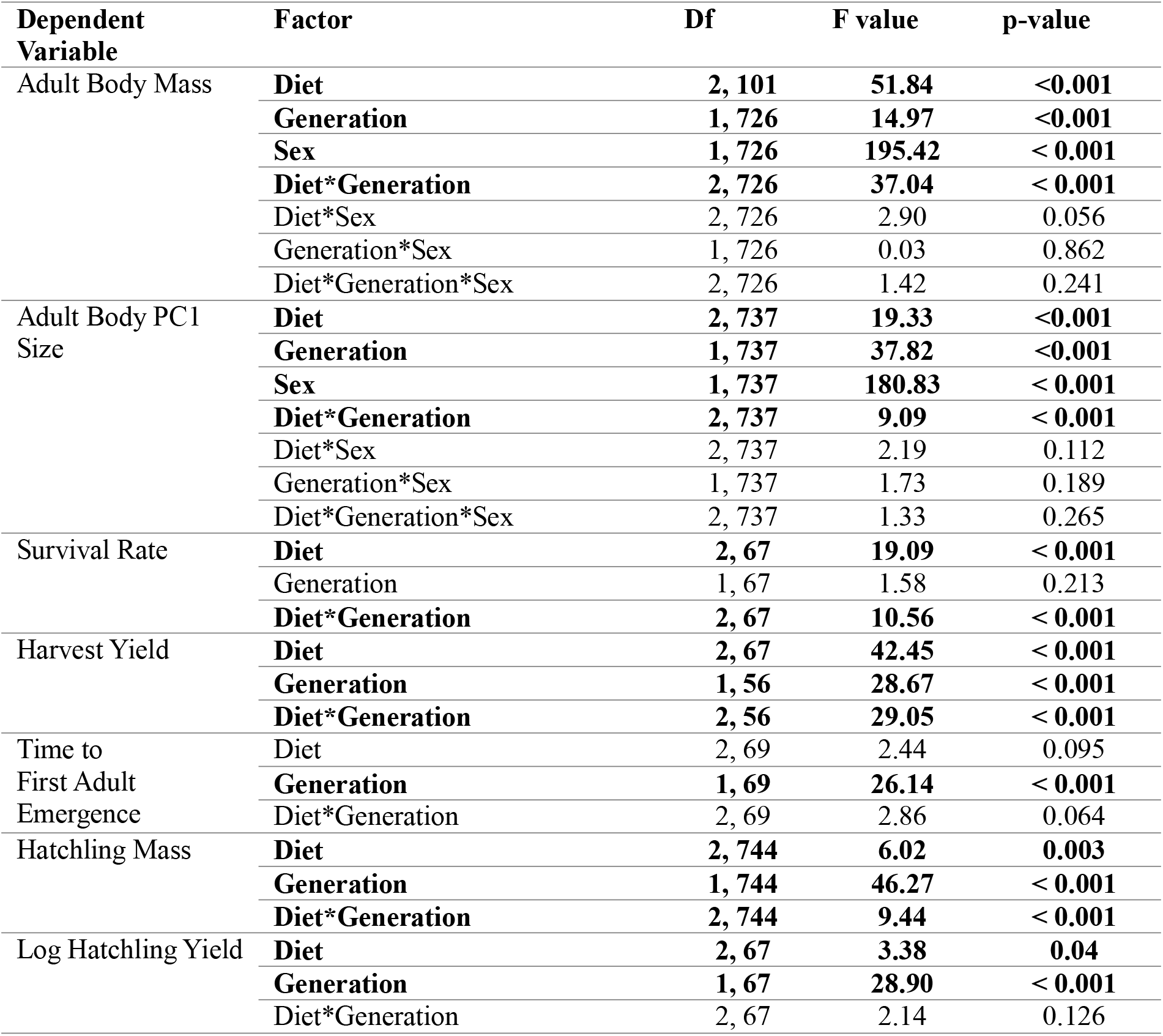
Analysis output of linear mixed effects models for quantified life history traits. Bolded values denote significance (p < 0.05).

## Results

### Adult Cricket Body Mass and Size

Diet treatment significantly impacted adult body mass (F_2,101_ = 51.84, p < 0.001, Figure 1a) and adult body size (F_2,737_ = 19.33, p < 0.001, Figure 1b) of *G. sigillatus*. On average, crickets reared on the high inclusion spent grain diet were 18.5% lighter (220 ± 4 mg) than control crickets (270 ± 5 mg) and this relationship remained consistent across five generations. Diet and generation interacted to influence both adult body mass (F_2,726_ = 37.04, p < 0.001) and adult body size (F_2,737_ = 9.09, p < 0.001). This interaction was driven by generational changes that occurred in crickets fed the gradual inclusion spent grain diet. The adult body mass of crickets reared on the gradual inclusion spent grain diet (288 ± 11 mg) did not differ from control crickets (273 ± 10 mg) in the first generation (15% spent grain), but a progressive decline in body mass was observed as the spent grain proportion was increased over successive generations. By generation five (75% spent grain) the crickets were 19.9% lighter (225 ± 10 mg) than control crickets (281 ± 11 mg), mirroring the stable effect of the 75% spent grain diet on body mass throughout the study. Sex explained a significant amount of the variation for both adult body mass (F_1,726_ = 195.42, p < 0.001) and adult body size (F_1,737_ = 180.83, p < 0.001). Generation, a variable used to account for fluctuations in environmental greenhouse conditions between generations, also explained a significant amount of the variation for both adult body mass (F_1,726_ = 14.97, p < 0.001) and adult body size (F_1,737_ = 37.82, p < 0.001), and was mainly driven by an unintended temperature fluctuation in generation three. Overall, our findings reveal that while cricket growth was strongly influenced by the proportion of spent grain in the diet, growth remained stable on a high inclusion diet despite a long-term change in diet.

**Figure 1.**
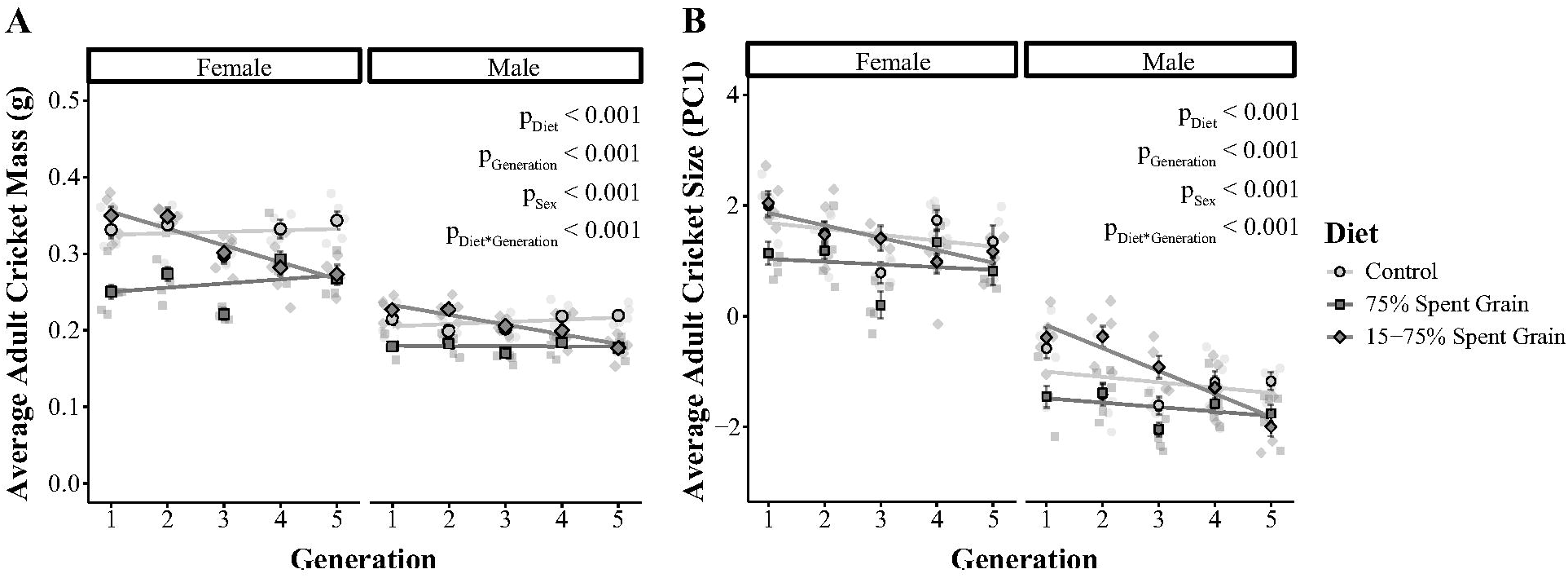
A) Adult mass (g) and B) body size (PC1, first principal component of morphological traits) across five generation of crickets reared on a control feed (standard farm feed, circles), a 75% spent grain feed (squares) or a feed that increased incrementally from 15-75% spent grain with each generation (diamonds). Each semi-transparent data point represents a replicate cricket colony. Solid points represent mean ± standard error.

### Survival and Development

Diet significantly impacted colony survival rates (F_2,67_ = 19.09, p < 0.001, Figure 2a). Cricket survival was similar between the high inclusion spent grain diet and the control for the last three generations, and about 9% higher on the high inclusion spent grain diet compared to the control in the first two generations. Crickets reared on the gradual inclusion spent grain diet had a 29.7% higher survival rate (85.8 ± 2.3 %) in the first generation compared to the control (60.3 ± 5.5 %). This positive impact on survival gradually declined over the five generations as inclusion of the spent grain diet increased. In the fifth generation, the survival rate on all diets was equal. Therefore, spent grain inclusion in the diet led to an increase in survival rate predominantly at low inclusion levels (15%). The time required for the first crickets to emerge to adulthood was similar across diets (F_2,69_ = 2.44, p = 0.095, Figure 2b). There was a significant effect of generation on time to adulthood (F_1,69_ = 26.14, p < 0.001, Figure 2b), which appeared to be driven mainly by the temperature fluctuation in generation three.

**Figure 2.**
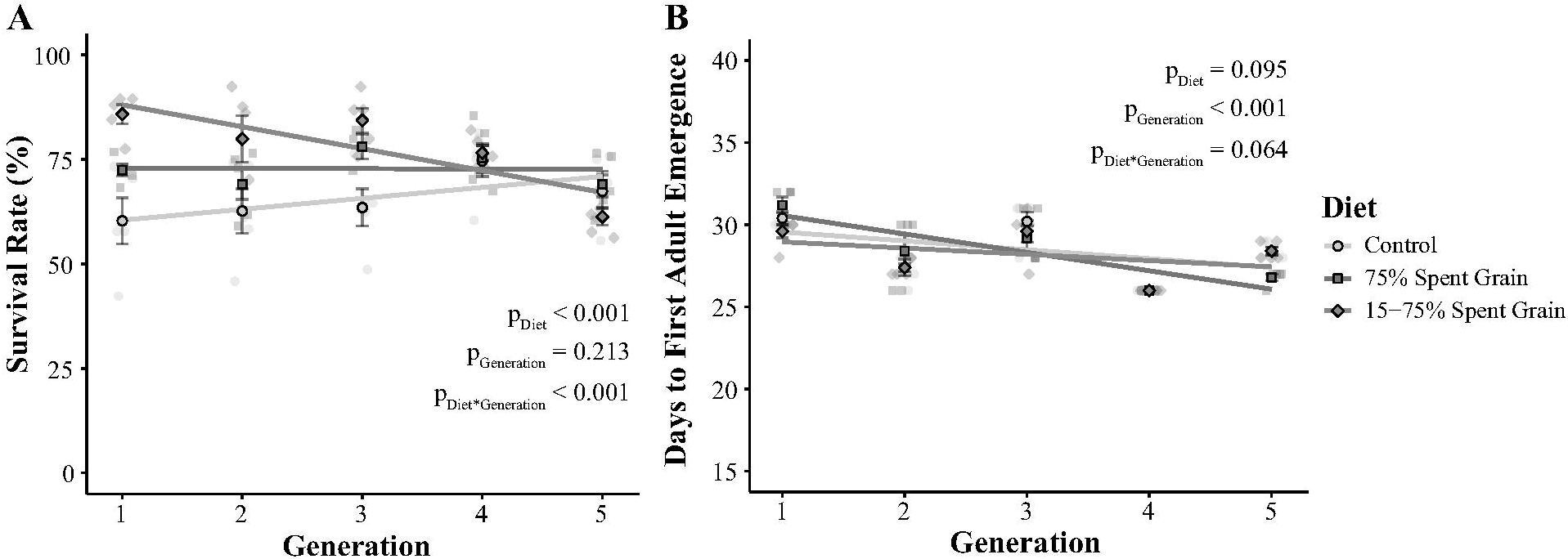
A) Survival rate (%) and B) time to first adult emergence (days) across five generation of crickets reared on a control feed (standard farm feed, circles), a 75% spent grain feed (squares) or a feed that increased incrementally from 15-75% spent grain with each generation (diamonds). Each semi-transparent data point represents a replicate cricket colony. Solid points represent mean ± standard error.

### Yield

Harvest yield, measured as the total weight of all crickets in the bin at the end of each generation, was significantly impacted by diet treatment (F_2,67_ = 42.45, p < 0.001, Figure 3). The high inclusion spent grain diet had a harvest yield that was 14.1% less (21.9 ± 4 g) than crickets reared on the control diet (25.5 ± 7 g). Diet and generation interacted to influence harvest yield (F_2,56_ = 29.05, p < 0.001); harvest yield on the gradual inclusion spent grain was 28.8% higher than on the control treatment in the first generation (34.4 ± 2 g), but gradually declined with successive generations. By the fifth generation, harvest yields did not differ between the high inclusion and the gradual inclusion spent grain diets (21.9 ± 4 g vs 19.1 ± 1 g). Increasing the proportion of spent grain in the cricket diet therefore led to decreased harvest yield over successive generations, but never did worse than the immediate effects of a high-inclusion spent grain diet. Low amounts of spent grain (15%), by contrast, significantly increased yield.

**Figure 3.**
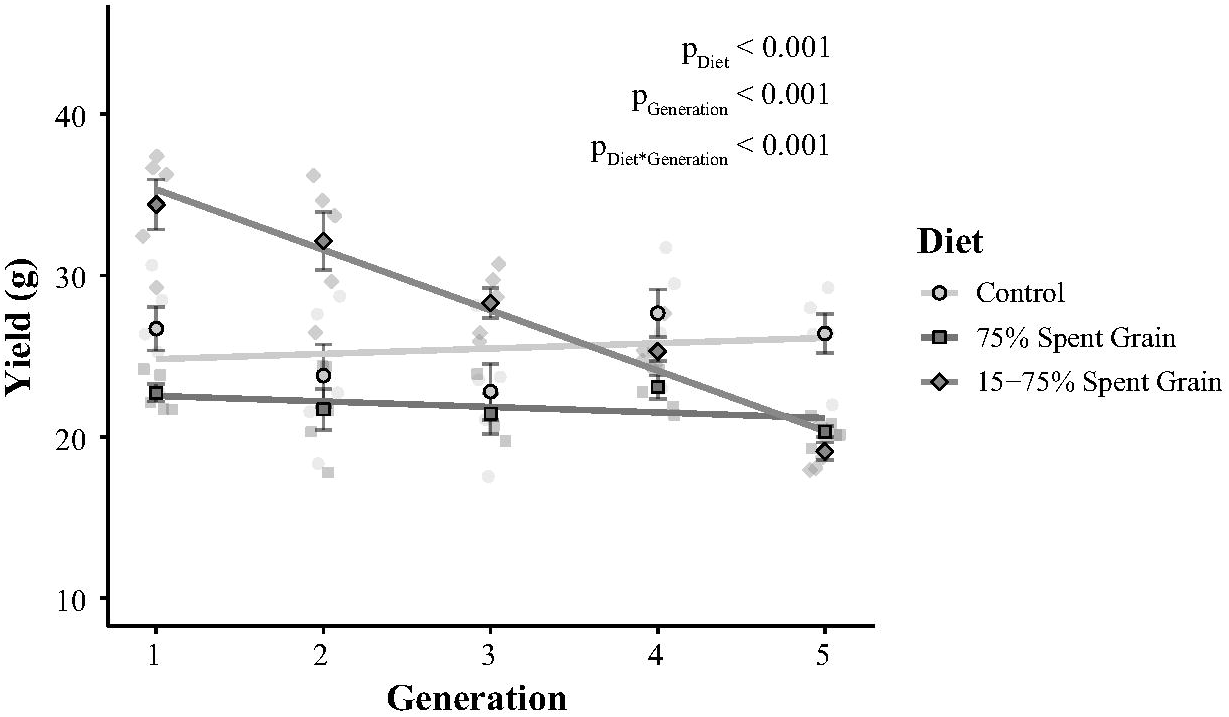
A) Yield (in g of crickets) across five generation of crickets reared on a control feed (standard farm feed, circles), a 75% spent grain feed (squares) or a feed that increased incrementally from 15-75% spent grain with each generation (diamonds). Each semi-transparent data point represents a replicate cricket colony. Solid points represent mean ± standard error.

### Hatchling Mass and Hatchling Yield

Diet had a significant effect on hatchling mass (F_2,744_ = 6.02, p = 0.003, Figure 4a) and hatchling yield (F_2,67_ =3.39, p = 0.04, Figure 4b). Hatchling mass displayed a variable trend, with hatchlings on the high inclusion (75%) spent grain diet being 11.9% smaller (0.89 ± 0.003 mg) compared to the control (1.01 ± 0.003 mg) in the first generation, but 8.2% larger and 21.3% larger than the control in the second and fifth generation, respectively. Crickets reared on the high inclusion (75%) spent grain diet produced approximately 40% fewer hatchlings (2.78 ± 0.08 g) in the first generation compared to the control (4.85 ± 0.32 g) and gradual inclusion spent grain (4.47 ± 0.74 g) diets, but by the fifth generation the mass of hatchlings produced was equal across all three diets. However, this change was not due to an increase in hatchlings produced on the high inclusion (75%) spent grain diet, but due to a decrease in hatchling yield on the control diet possibly from the temperature fluctuation in generation three. Therefore, hatchling yield, but not mass, was consistently reduced by a high spent grain content in the diet.

**Figure 4.**
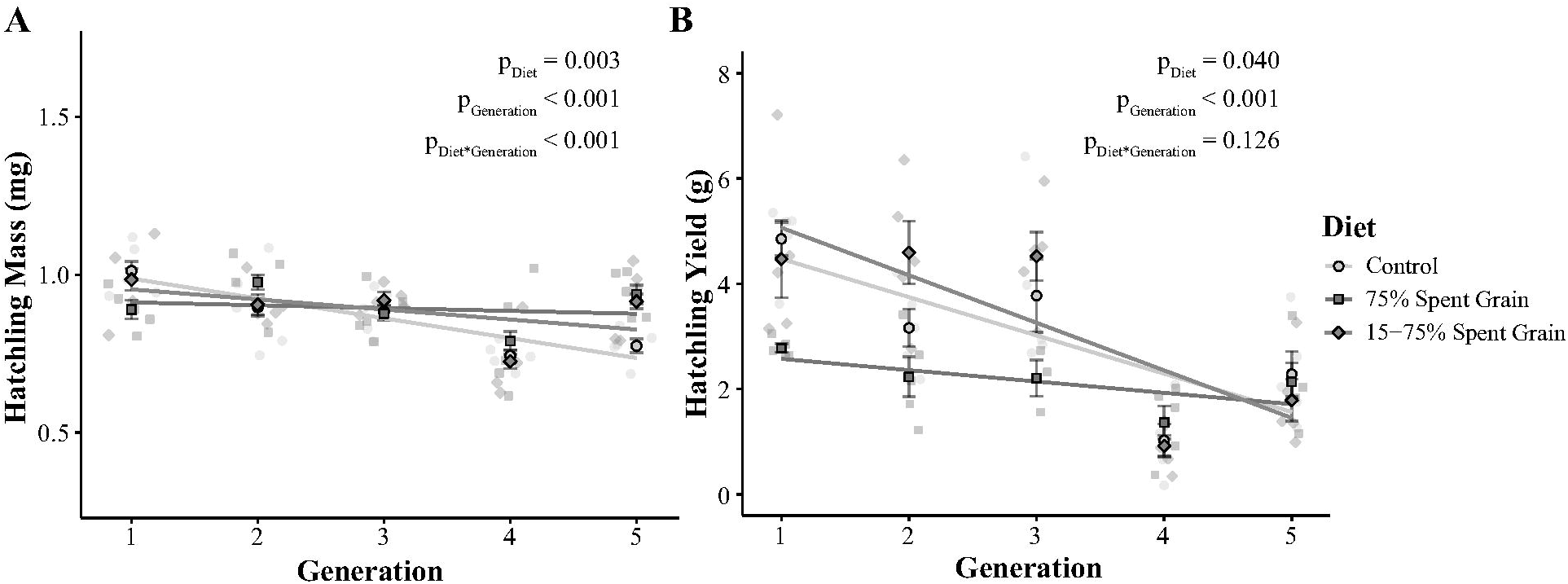
A) Individual hatchling mass (g) and total yield of hatchlings (g) across five generation of crickets reared on a control feed (standard farm feed, circles), a 75% spent grain feed (squares) or a feed that increased incrementally from 15-75% spent grain with each generation (diamonds). Each semi-transparent data point represents a replicate cricket colony. Solid points represent mean ± standard error.

## Discussion

Our results are the first to demonstrate that brewer’s spent grain is a suitable bulk feed ingredient for long-term rearing of commercially produced crickets. We found that adding spent grain to diets has minimal long-term effects on key life history traits important to production of farmed crickets, and that relative performance remains consistent over successive generations. This is promising from a farming perspective, as the consistent performance experienced by crickets raised on a high spent grain diet indicates that cheaper waste-based diets have potential to work as long-term feed without the risk of colony collapse. By furthering knowledge on the short- and long-term impacts of waste diets on cricket life history traits, our results provide insight into how insects respond to long-term dietary shifts in addition to better informing producers on integration of waste products into commercial insect feed. These findings have important implications for both the edible insect industry and meeting global waste-reduction goals, particularly as low inclusion spent grain diets have the potential to repurpose organic waste to both reduce feed costs and improve farm productivity.

These results are biologically intriguing, as the lack of multigenerational change provides insight into the relationship between parental conditions and offspring performance. Multiple hypotheses have been proposed on how environmental stressors like suboptimal nutrition in the parental generation may impact offspring fitness through non-genetically transmitted parental effects. For example, a parental stress hypothesis predicts that poor nutrition experienced by the parents may lead to poorer quality offspring (Frago and Bauce, 2014; Vijendravarma *et al*., 2010). Conversely, an adaptive hypothesis predicts that parents reared on a poor diet may be able to better prepare their offspring to better tolerate the same conditions (Badyaev and Uller, 2009; Burgess and Marshall, 2014; Mousseau and Fox, 1998). Both scenarios incorporate transgenerational effects, where the phenotype of the parent influences the phenotype of the offspring (Mousseau and Fox, 1998; Shikano *et al*., 2015). While our experimental design did not directly test for parental effects, our results did not show evidence of a changing response to diet over time despite phenotypic variation (e.g., reduced growth) on a high spent grain diet observed in a single generation. These findings support neither an adaptive nor maladaptive transgenerational effects hypothesis, thus raising questions about how insects like crickets remain resilient to nutritional challenges over the long-term.

As demonstrated in single-generation studies, crickets are very tolerant of a wide-range of dietary nutritional profiles (Muzzatti *et al*., 2024). This tolerance may help buffer crickets against transgenerational effects associated with poor parental nutrition. This dietary flexibility is supported by plasticity in behaviour and digestive physiology that allows insects to extract essential nutrients from a variety of diet components. For example, insects may behaviourally compensate for dietary imbalance by regulating their feed intake, particularly with respect to protein (Raubenheimer, 1992). In our study, crickets raised on a high inclusion (75%) spent grain diet required an increased volume of feed to permit *ab libitum* feeding (Figure S1), supporting the hypothesis that they may be partially adjusting for poor nutrition by consuming more. In addition, physiological mechanisms like regulation of digestive enzyme secretion in the gut can help balance nutrient assimilation (Lazarević *et al*., 2023; Woodring and Weidlich, 2016). Black soldier fly larvae fed a nutritionally unbalanced diet mimicking fruit and vegetable waste displayed differences in digestive enzyme activity in addition to energy reserves and midgut cell morphology (Bonelli *et al*., 2020) These results were correlated with only moderate impacts of diet on life history (Bonelli *et al*., 2020), indicating that gut plasticity likely plays a role in compensation for poor nutrition in generalist insects. Future studies exploring digestive markers like enzyme profiles of crickets reared on spent grain or other waste-based diets could help provide insights into the mechanisms of long-term dietary tolerance.

Despite high survival and the completion of successful life cycles across all diets, high inclusion of spent grain reduced adult body mass by about 18.5%, leading to reductions in overall harvest yield, and decreased hatchling output by about 40%. A likely contributor to these changes is the high content of undigestible lignocellulosic material (i.e., fiber) in spent grain (Aliyu and Bala, 2011; Ikram *et al*., 2017). Fiber is not readily broken down by endogenous enzymes into accessible nutrients and has been linked to nutrient dilution and decreased nutrient absorption in pigs (Bach Knudsen, 2001) and poultry (Tejeda and Kim, 2021). Development and reproduction are energetically costly processes, and access to dietary protein and digestible carbohydrates in the diet is critical for optimal insect growth (Clark *et al*., 2015; Harrison *et al*., 2014; Muzzatti *et al*., 2024) and fecundity (Gutiérrez *et al*., 2020; Magara *et al*., 2019; Rapkin *et al*., 2018; Rho and Lee, 2016). Therefore, a more limited availability of key nutrients to allocate towards growth and reproduction despite compensatory actions likely led to declines in harvest yield for spent grain-fed crickets. Our results align with past (single generation) studies on crickets reared on organic waste-containing diets. For example, *Acheta domesticus* reared on reared on diets in which soy was replaced with spent grain (∼ 41% of total diet) were slightly smaller than crickets fed a control feed (Sorjonen *et al*., 2019). Similarly, *Acheta domesticus* reared on mixed waste diets grew smaller than those reared on a control diet, which was linked to a lower nitrogen (protein) to fiber ratio (Lundy and Parrella, 2015). While reproductive metrics are less explored in studies on waste diets for farmed crickets, poor nutrition is often linked to decreased maternal investment in insects. For example, crickets (*Scapsipedus icipe*) raised on balanced diets had higher lifetime fecundity than crickets raised on protein-poor diets (Magara *et al*., 2019). As hatchlings produced on the high spent grain diet were viable and showed no evidence of increased mortality, females may have been prioritizing offspring fitness over offspring number to compensate for limited nutrition. Adjusting investment per offspring to produce fewer, better provisioned offspring is an accepted life history strategy (Smith and Fretwell, 1974). Egg size is positively correlated to fitness in species lacking maternal care (Azevedo *et al*., 1997), and increases in egg size produced by insects reared in poor diets despite reduced body size has been documented in species like *Drosophila* (Vijendravarma *et al*., 2010). This prioritization could potentially explain why crickets produced larger hatchlings on the high inclusion (75%) spent grain diet relative to the control in generation five. Taken together, our findings suggest that while crickets can tolerate high-spent grain diets, nutritional limitations, likely related to fiber content, constrain growth and reproductive output. In particular, the large impact on hatchling output indicates that high inclusion spent grain diets may be better purposed as feed for production animals, rather than feed for breeding stock.

In the event that adaptive transgenerational effects were observed leading to increased cricket performance on the high inclusion (75%) spent grain diet in the fifth generation compared to the first, we included a gradual inclusion diet to evaluate whether the initial negative impacts could be mitigated using a “weaning” approach. However, cricket growth on this diet linearly decreased with increasing proportions of spent grain over the course of the five generations, further supporting the idea that spent grain inclusion influences cricket performance, but that this relationship is not being strongly impacted by parental diet through transgenerational effects. Interestingly, the lower inclusion (15-30%) diets in generations one and two led to nearly a 30% increase in survival and yield. It may be the case that while high levels of dietary fiber lead to nutrient dilution, lower levels increase performance through beneficial actions like promotion of healthy gut microbial communities (Douglas, 2015; Engel and Moran, 2013). These results suggest that spent grain is a valuable feed ingredient that has the capacity to both reduce costs and increase productivity of cricket farming when used in moderation, reinforcing the role of cricket in food waste reduction and circular bioeconomy initiatives.

Beyond diet, unintentional variations in environmental conditions influenced cricket life history. A reduction in body mass was observed on all three diets in the third generation, as well as a reduction in hatchling yield on the control in the fourth and fifth generations. Although greenhouse conditions were relatively stable throughout the ten-month experiment, slight fluctuations were accounted for in all models as the ‘Generation’ factor. This factor explained a significant amount of variation for most measured traits including the control. In the third generation, a relatively modest drop in temperature from a daily average of approximately 30.5°C to 27 °C was experienced by crickets for roughly ten days due to a heating system malfunction (Figure S2). Environmental factors like temperature, humidity and light cycles are known to strongly impact insect life history traits (Johnsen *et al*., 2021; Ratte, 1984), and may also interact with diet (Clissold *et al*., 2013; Kingsolver and Woods, 1998; Kutz *et al*., 2019). In *Gryllodes sigillatus* crickets, rearing temperatures between 30-36 °C are considered optimal, with lower temperatures negatively impacting performance (Kong *et al*., 2025). Therefore, the fluctuations in insect growth and fecundity observed are likely tied to sub-optimal temperatures during development. Interestingly, the impacts on fecundity were not tied directly to the generation exposed to the temperature drop (generation three), but persisted in the control group into generations four and five. Transgenerational effects of thermal stress in insects remains poorly understood, but both low and high temperature shock have been shown to induce effects on fecundity that last for two generations in fall armyworm (Reshma *et al*., 2023). This suggests that even short-term temperature stressors have profound consequences for insect performance in the long-term, highlighting the need for studies investigating transgenerational plasticity of insects in response to temperature.

Overall, the results of our study demonstrate the capacity for crickets to survive on diets containing spent grain for multiple generations, indicating that spent grain is a promising ingredient for inclusion in cricket feed that could be used as a cost-effective alternative to traditional feed components. While high proportions of spent grain in the diet are nutritionally challenging for crickets and lead to reduced growth compared to a control, crickets demonstrate a remarkable ability to survive and grow on waste-based diets for prolonged periods of time without compounding adverse effects. Replacement of 75% of traditional feed with spent grain only led to a 14% drop in yield relative to the control, while replacement of lower proportions (15-30%) led to up to a 29% increase in yield. This indicates that, in addition to being cost-effective, partial inclusion of spent grain into farm feeds may even increase yield while decreasing production cost.

## Acknowledgements

Our thanks to the individuals at Entomo Farms, Stray Dog Brewing Company and Aspire Food Group for their continued support and contributions to this work.

## Author Contributions

Experiment conceptualization and design were performed by S.Y.K., S.M.B, and H.A.M. Rearing experiments was performed by S.Y.K. Supervision was performed by S.M.B. and H.A.M. Project administration, funding acquisition and resource provision were performed by S.M.B and H.A.M. Data visualization and analysis was performed by S.Y.K. Original draft was written by S.Y.K. All authors contributed to editing and review.

