## Supplemental Tables & Figures for "Farmed cricket performance remains stable over five generations of rearing on a waste-based diet"

Kasdorf, S.Y., Bertram, S.M. and MacMillan, H.A.

**Supplemental Tables & Figures**

**Table S1.** Diet provision for each treatment group by generation

| Diet | Generation | | | | |
| --- | --- | --- | --- | --- | --- |
|  | 1 | 2 | 3 | 4 | 5 |
| Control | Farm Feed | Farm Feed | Farm Feed | Farm Feed | Farm Feed |
| High Inclusion Spent Grain | 25% Farm Feed,  75% Spent Grain | 25% Farm Feed, 75% Spent Grain | 25% Farm Feed, 75% Spent Grain | 25% Farm Feed, 75% Spent Grain | 25% Farm Feed, 75% Spent Grain |
| Gradual Inclusion Spent Grain | 15% Farm Feed, 75% Spent Grain | 30% Farm Feed, 75% Spent Grain | 45% Farm Feed, 75% Spent Grain | 60% Farm Feed, 75% Spent Grain | 75% Farm Feed, 75% Spent Grain |

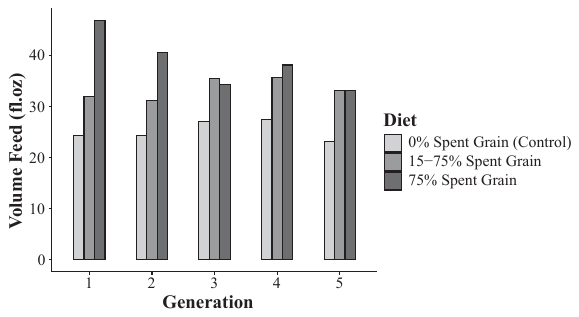

**Figure S1.** Total volume of feed in fluid ounces provided to crickets per generation on a control feed (standard farm feed, light grey), a 75% spent grain feed (medium grey) or a feed that increased incrementally from 15-75% spent grain with each generation (dark grey) to maintain *ab libitum* feeding.

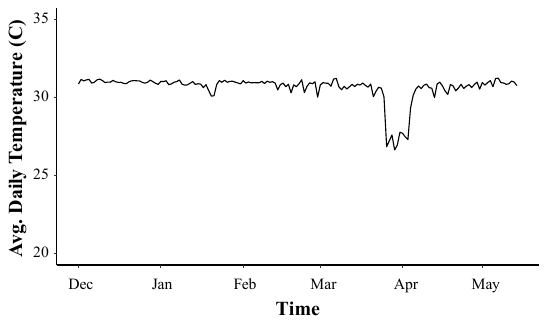

**Figure S2.** Average daily temperature (°C) of the greenhouse used to house crickets during the multigenerational experiment. The data shown is from 2023-12-01 (YYYY-MM-DD) to 2024-05-15 (YYYY-MM-DD). The experiment ran from 2023-12-01 (YYYY-MM-DD) to 2024-08-11 (YYYY-MM-DD). A drop in average daily temperature is visible for a period of time at the end of March.
